# Fibroadipogenic Progenitors contribute to microvascular repair during skeletal muscle regeneration

**DOI:** 10.1101/2021.09.06.459112

**Authors:** David Ollitrault, Valentina Buffa, Rosamaria Correra, Angeliqua Sayed, Bénédicte Hoareau, Sophie Pöhle-Kronawitter, Sigmar Stricker, Jean-Sebastien Hulot, Mariana Valente, Giovanna Marazzi, David Sassoon

## Abstract

Skeletal muscle injury results in a disruption of the muscle bed vascular network. A local source of vascular progenitors during muscle regeneration has not been clearly identified. Fibroadipogenic progenitors (FAPs) are required for proper regeneration, however they can also directly contribute to fibrotic and fatty infiltration in response to chronic muscle injury and muscle disease. We show here that acute muscle injury leads to hypoxia and glucose deprivation, triggering FAP proliferation and differentiation into endothelial cells *in vitro* and *in vivo*. In response to glucose deprivation, FAPs down regulate fibrotic and fat associated genes and acquire an endothelial cell fate, which is dependent upon mTORC2-HIF2α-eNOS pathway. These findings bring new insights into the mechanisms of vascular regeneration during muscle regeneration and define a highly plastic resident progenitor population that responds to oxygen/glucose-deprivation induced cell stress by promoting an endothelial cell fate.

## Introduction

Acute skeletal muscle injury leads to the degeneration of myofibers as well as the associated vascular network. Vascular tissue is required for oxygen and glucose delivery, consequently vascular regeneration is a critical step towards muscle repair. Several muscle tissue resident progenitors have been identified, most notably satellite cells that generate new myofibers, as well as the fibro-adipogenic progenitors (FAPs) that give rise to fat and connective tissue (*1–5*). FAPs also secrete key cytokines in response to muscle injury that support satellite cell expansion and differentiation (*4, 6–9*). We identified previously a population of interstitial cells based upon the expression of the pan-stem cell marker, *Pw1/Peg3* (termed PICs for PW1+ interstitial cells) (*1, 2*). PICs consist largely of PDGFRα +/CD34+/SCA1+/Lin- cells, which comprise the entire FAP population, as well as PDGFRα-/CD34+/SCA1+/Lin- cells, which have multiple cell fate potentials including smooth muscle, skeletal muscle, fat, and fibroblasts (*3, 10*). It has been recently shown that FAPs can also give rise to ectopic bone in muscle tissue (*11, 12*) and participate in osteo-chondrogenesis during bone repair (*13*), demonstrating a remarkable cell fate plasticity.

Vasculogenesis is stimulated by the release of cytokines such as Vascular Endothelial Growth Factor (VEGF), however signals that initiate endothelial cell (EC) differentiation are less characterized (*14*). It has been shown that hypoxia and/or nutrient deprivation can regulate angiogenesis (*15–19*), leading us to speculate whether these signals are present following acute muscle injury and if they contribute to subsequent vascular repair. Here, we demonstrate that acute muscle injury results in a hypoxic state, triggering a HIF-1α dependent proliferative response in FAPs. Following proliferation, FAPs respond to glucose deprivation by progression towards endothelial differentiation through a HIF-2α-eNOS dependent pathway. Using lineage tracing, we demonstrate that FAPs contribute to microvasculature formation as well as self-renewal following injury *in vivo*.

## Results

### Hypoxia induces FAP proliferation

Muscle injury leads to localized tissue hypoxia due to the loss of vascularity in the damaged area (*20–23*). To assess hypoxia following muscle injury, we examined the *tibialis anterior* muscle (TA) of mice injected with pimonidazole, which accumulates in hypoxic cells(*24*), following cardiotoxin (CTX) induced injury. Uninjured muscle exhibited a weak and punctate staining, whereas a strong and uniform staining was detected 5 days after CTX injection (Fig. 1A). Consistent with previous reports (*4, 25*), we observed that FAPs underwent a marked expansion in cell number following injury (Fig. 1B), indicating that they can proliferate under hypoxic conditions. Cell expansion under hypoxic conditions is typically mediated by the upregulation and nuclear translocation of the Hypoxia-Inducible Factor-1 alpha (HIF-1α) (*26, 27*). In uninjured muscle, HIF-1α is detected primarily in the cytoplasm of interstitial cells (Fig. 1C), whereas it is located in the nuclei of interstitial cells and newly regenerated myofibers 5 days post-injury (5 dpi, Fig. 1C). To determine whether HIF-1α expression is enriched in FAPs, we FACS-sorted FAPs and satellite cells as described previously (Fig. S1) (*2*). We observed high levels of *HIF-1α* transcript and protein in FAPs as compared to satellite cells (Fig. 1D, E).

Consistent with the localization of HIF-1α expression in interstitial cells *in vivo*, we observed cytoplasmic HIF-1α in freshly sorted FAPs (Fig. 1E). We next tested whether a decrease in oxygen levels induces proliferation in HIF-1α expressing FAPs *in vitro*. When FAPs are grown under hypoxic conditions (1.5% O_2_), we observed HIF-1α nuclear translocation in FAPs, whereas purified satellite cells showed no nuclear localization (Fig. 1F). Furthermore, we observed an increase in the proliferation of FAPs in response to hypoxia (Fig. 1G). Previous studies have shown that the mammalian target of rapamycin complex 1 (mTORC1) activates HIF-1α translation, synthesis and translocation (*28*). We observed that the mTORC1 inhibitor, rapamycin (*28*), markedly reduced FAP proliferation under hypoxia and inhibited HIF-1α nuclear translocation (Fig. 1H, I). Furthermore, siRNA-mediated gene silencing of *HIF1-α* abrogated hypoxia-induced FAPs proliferation (Fig. 1J and Fig. S2). Taken together, these data suggest that the mTORC1-HIF-1α pathway regulates hypoxia-induced FAP proliferation. As both rapamycin and HIF-1α silencing caused a modest reduction in cell proliferation under normoxic conditions, we propose that this pathway is active regardless of oxygen levels as has been reported for other cell types (*29*).

**Fig. 1.**
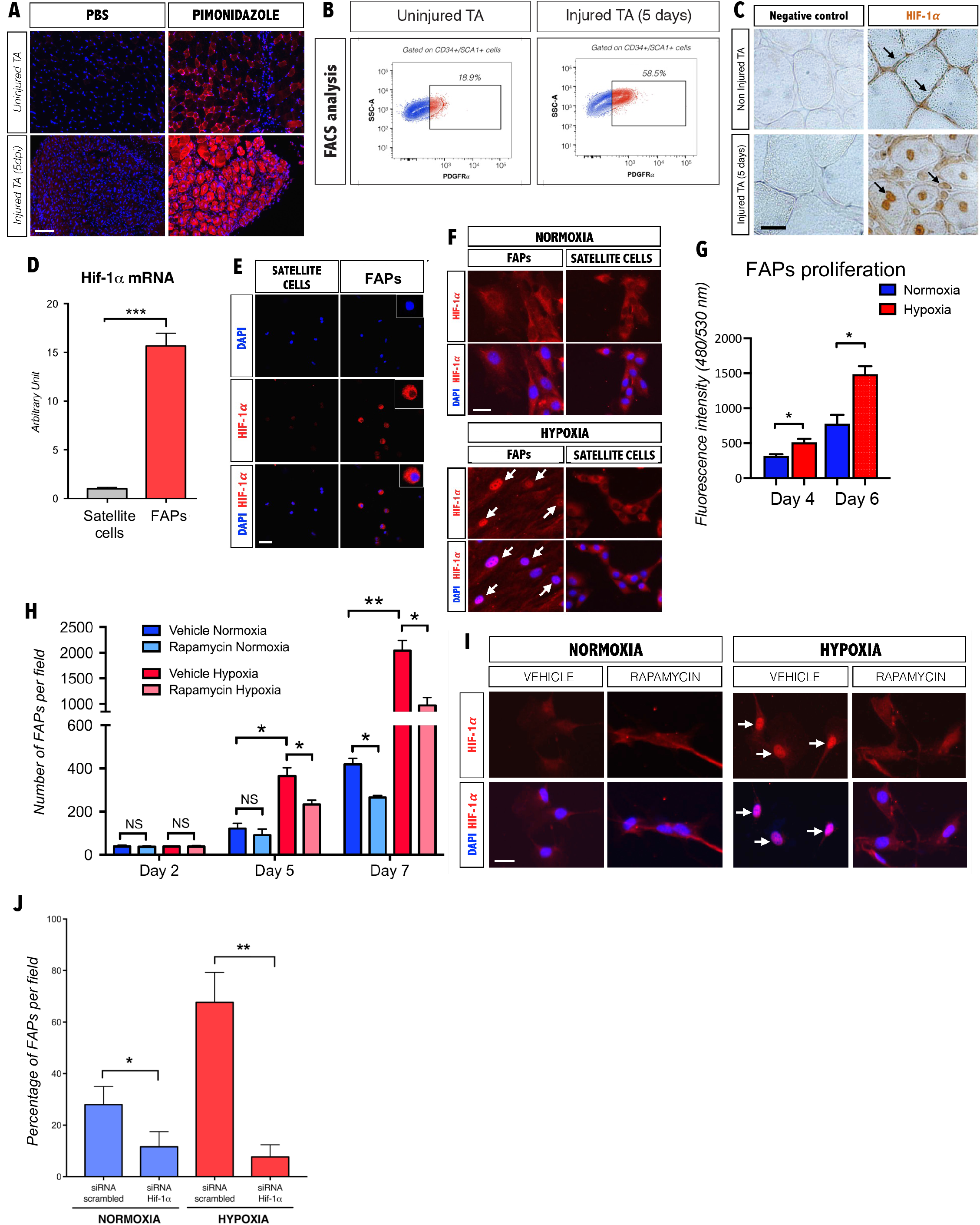
Muscle injury results in hypoxia that triggers FAPs proliferation through the mTORC-HIF-1α pathway. **(A)** Representative photomicrographs of pimonidazole immunostaining (hypoxia marker) of cross sections of the TA 5 days following CTX-injury (5 dpi) as compared to uninjured muscle. Nuclei stained with DAPI. Scale bar, 100μm (n=3). **(B)** Representative FACS analyses showing an increase in the percentage of PDGFRα+ cells 5 days following injury (n=3). **(C)** HRP immunostaining of HIF-1α of uninjured and 5 days post injured TA. Scale bar, 20μM (n=3). **(D)** Histogram showing qPCR levels of HIF-1α from freshly FACS-sorted satellite cells and FAPs (PDGFRα+ cells) (n=3). **(E)** HIF-1α and DAPI staining of freshly FACS-sorted cytospun satellite cells and FAPs (PDGFRα+ cells). Scale bar, 25μM. Inset panels showing higher magnification to reveal predominantly cytoplasmic staining of HIF-1α (n=3). **(F)** Representative HIF-1α immunostaining of FAPs (PDGFRα+ cells) and satellite cells cultured 24 hours under normoxic and hypoxic conditions. Nuclei were counterstained with DAPI. Nuclear translocation is detected only in FAPs (arrows) Scale bar, 20μm (n=3). **(G)** Cell proliferation assay of FAPs (PDGFRα+ cells). FAPs were cultured 4 or 6 days in normoxia and hypoxia (n=4). **(H)** Time course analysis of FAPs (PDGFRα+ cells) proliferation under normoxic and hypoxic conditions with or without 100nM of rapamycin (mTORC1 inhibitor). Values represent the mean values of number of cells per fields from 3 independent experiments (n=3). **(I)** HIF-1α immunostaining of PDGFRα+ cells cultured under normoxia or hypoxia for 5 days with or without 100nM rapamycin (arrows denote nuclear translocation). Nuclei were counterstained with DAPI. Scale bar, 20μm (n=3). **(J)** Mean percentage value of PDGFRα+ cells cultured 72 hours in normoxia and hypoxia following HIF-1α siRNA or scrambled siRNA (n=4). In all graphs values represent the mean ± s.e.m. \**P*<0.05, \*\**P*<0.01 and \*\*\**P*<0.001.

### Hypoxia increases glucose uptake in FAPs through the mTORC1-HIF-1α-Glut1 pathway

HIF-1α cell proliferation is typically coupled with an increase in cellular anaerobic glucose metabolism in response to hypoxia (*30*) and an increase in glucose metabolism can be measured by lactate production (*31*). As expected, we observed an increase in lactate production from FAPs grown under hypoxic conditions consistent with an increase in glucose uptake (Fig. 2A). Glucose uptake occurs through glucose transporters (GLUT) and is essential for anaerobic cell proliferation (*30, 32*). Previous studies have demonstrated that HIF-1α regulates GLUT1 expression and GLUT1 is the most ubiquitously expressed glucose transport molecule (*26*). We therefore confirmed *GLUT1* expression in FAPs and that *GLUT1* expression was upregulated in FAPs in response to hypoxia (Fig 2B) and blocked following *HIF-1α* silencing (Fig. 2B and Fig. S2). Pharmacological glucose transport inhibitors have been developed to target cancer cells that typically rely upon glycolysis for expansion (*33*). Using the GLUT1 inhibitor, Wzb-117 (*34*), we observed a marked reduction of FAP proliferation regardless of oxygen levels, whereas satellite cells were unaffected (Fig. 2C), demonstrating that FAP proliferation is primarily GLUT1-dependent, although it is likely other GLUT isoforms are present.

**Fig. 2.**
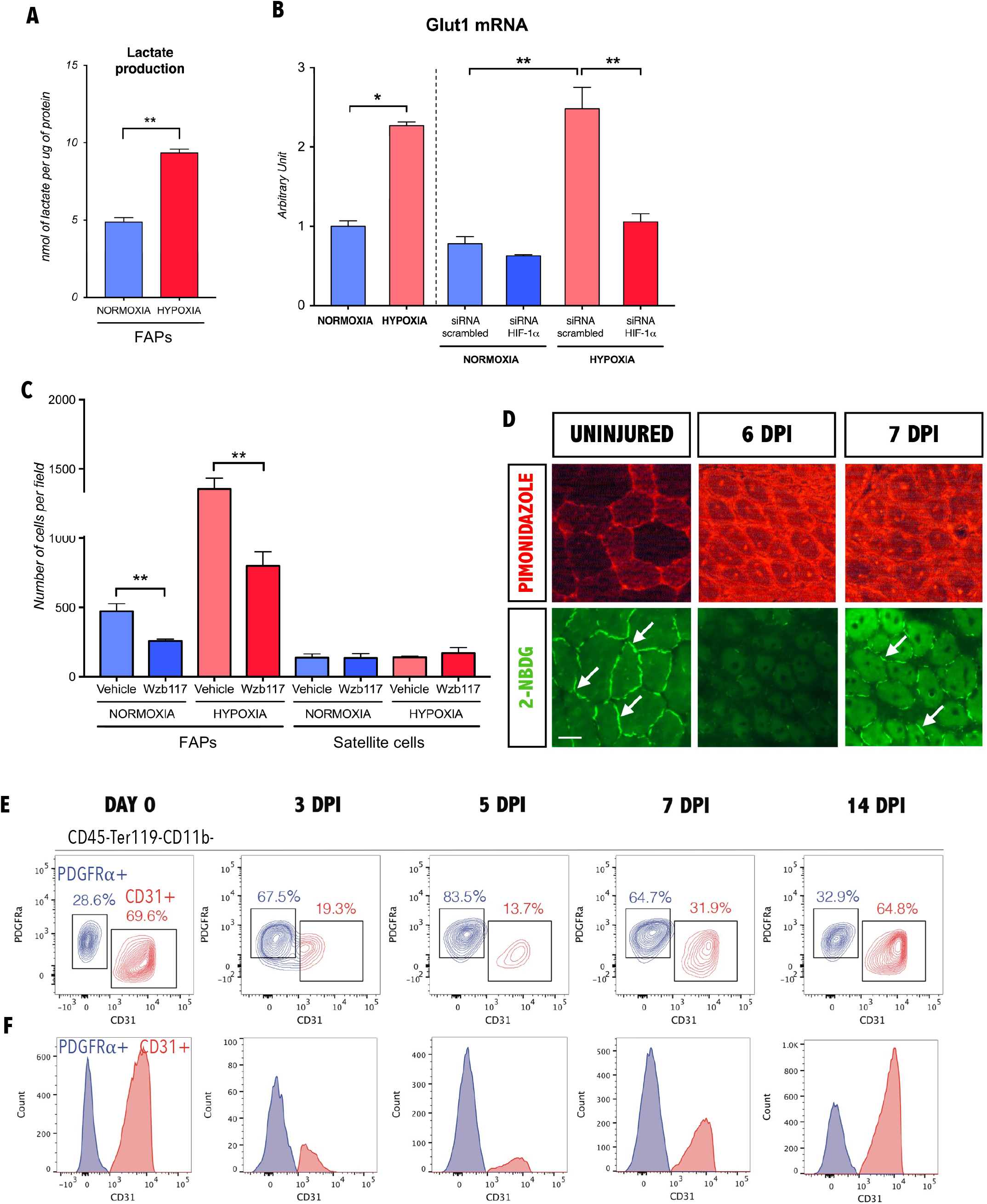
Inhibition of HIF-1α targets GLUT1 and abrogates FAPs proliferation. **(A)** Quantification of lactate production from FAPs (PDGFRα+ cells) cultured for 16 hours in DMEM (high glucose) without FBS. Supernatant lactate levels were measured normalized to protein content (n=3). **(B)** qPCR of GLUT1 from FAPs (PDGFRα+ cells) cultured under normoxia or hypoxia in response to HIF-1α iRNA knockdown (n=4). **(C)** Quantification of FAPs (PDGFRα+ cells) and satellite cells number cultured for 7 days under normoxia or hypoxia, with or without 10μM of GLUT1 inhibitor Wzb-117 (n=3). **(D)** Pimonidazole and 2-NBDG staining in TA muscle uninjured and 6 and 7 days following injury (dpi) (n=3). Scale bar, 50μM. **(E)** Time course FACS analysis of PDGFRα and CD31 expression during regeneration in skeletal muscle (3, 5, 7 and 14 days following CTX-injury from 3 month-old C57Bl/6 mice) and in uninjured muscle. Blue: PDGFRα+/CD31-cells, Red: CD31+ cells (n=3). **(F)** Counts of FACS analysis of PDGFRα+/CD31-(blue) and CD31+ cells (red) in uninjured muscle, 3, 5, 7 and 14 dpi (n=3). In all the graphs, values represent the mean ± s.e.m. \**P*<0.05, \*\**P*<0.01 and \*\*\**P*<0.001.

Given our observations that acute muscle injury leads to tissue hypoxia and that FAPs respond to both oxygen and glucose availability, we analyzed hypoxia and glucose uptake using pimonizadole and fluorescent labeled 2 deoxy D-glucose (2NBDG) (*35*), respectively, as well as the dynamic of FAP and EC compartments during the first week after injury (Fig. 2D). In uninjured muscle, 2NBDG is restricted to the interstitium (Fig. 2D), consistent with the observation that FAP proliferation is dependent upon glycolytic metabolism (Fig. 2C). However, while injured muscle remains hypoxic until 7 dpi (Fig. 2D), we observed a decrease in glucose accumulation in the interstitial space at 6 dpi, which recovers by 7 dpi (Fig. 2D). Parallel to the microenvironment alteration observed during the first week after injury, the FAP (PDGFRα+/CD31-cells) compartment showed an expansion of the population at 3 and 5 dpi, whereas the ECs (CD31+/ PDGFRα- cells) expansion emerged later around 7 and 14 dpi (Fig. E, F). Taken together, these data show that FAP population expanded in a hypoxic and glucose deprived environment during the first week of regeneration.

### Glucose and oxygen levels govern FAP cell fates

FAPs readily differentiate to adipocytes when grown under proadipogenic conditions (*2, 4*). We observed that high glucose levels are sufficient to promote adipogenesis in FAPs grown under normoxia, however adipogenesis was completely inhibited by hypoxia observed with the absence of cells stained for Oil Red O (Fig. 3A) and the decrease of the adipogenic transcription program (adiponectin (*Acr30)*, *lipoprotein lipase (Lpl), peroxisome proliferator activated receptor delta (Pparδ))* (Fig 3B). Additionally, our results using a GLUT1 inhibitor Wzb117 (Fig 2C) indicate that FAPs require glucose to proliferate in response to hypoxia, however it is likely that glucose transport is maintained in the presence of the inhibitor, albeit at lower levels, through other glucose transporters isoforms. It has been shown that the glucose analog, 2-deoxy-D-glucose (2-DG), completely blocks glycolysis in cultured cells (*36*), providing a tool to test directly the dependence of FAPs for glucose uptake in response to hypoxia. We therefore plated freshly FACS-sorted FAPs, as well as satellite cells and dermal skin fibroblasts that were allowed to reach 80% confluency followed by 48 hours treatment with 2-DG. Following 2-DG treatment, we observed pronounced cell death in satellite cells and no overt changes in skin fibroblasts (Fig. 3C, D). In contrast, FAPs acquired a branched morphology with the presence of meshes or branching points and elongated segments (tube) following 2-DG treatment (Fig. 3C).

**Fig. 3.**
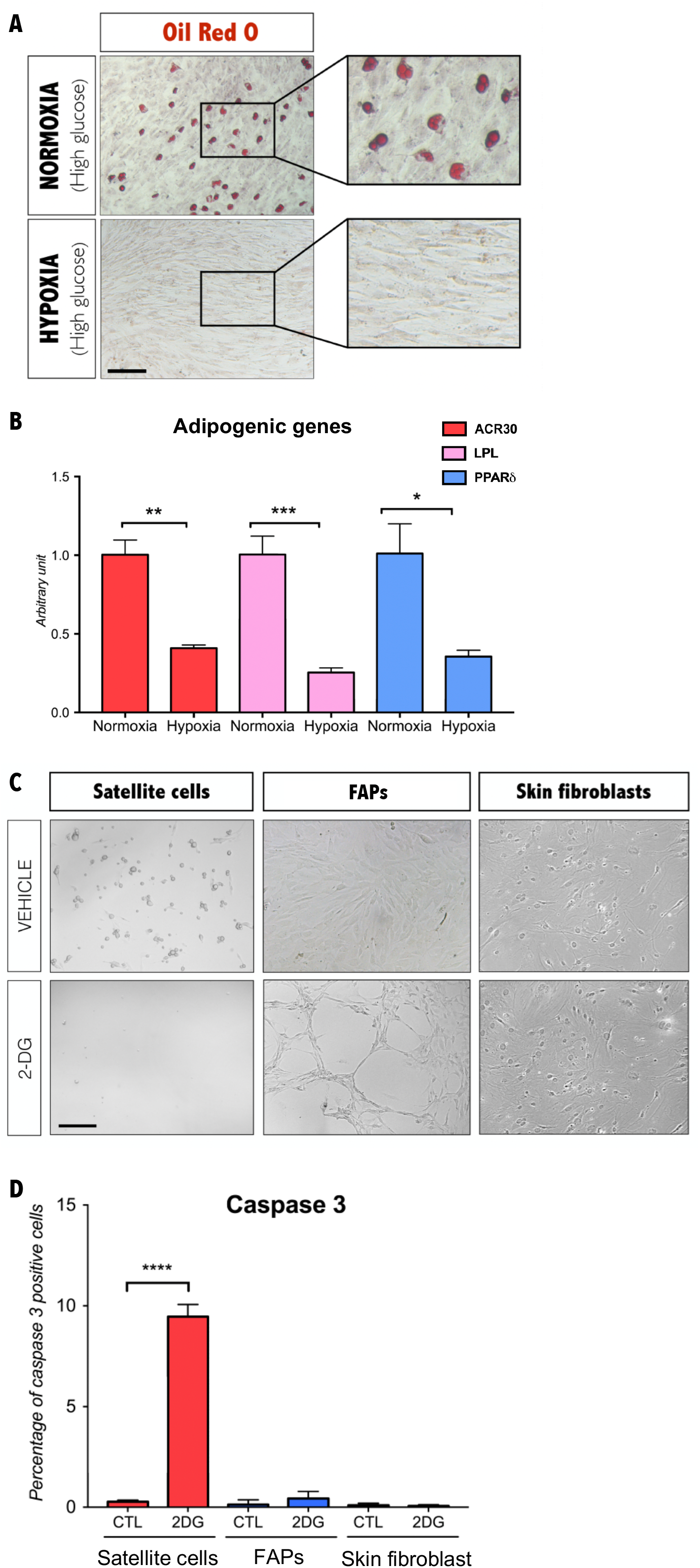
FAPs survive total glucose deprivation and hypoxia inhibits FAPs adipocyte cell fate. **(A)** Oil red O staining of FAPs cultured in high glucose in normoxia or hypoxia (n=3). Scale bar, 200μm. **(B)** qPCR of Acr30, Lpl, Ppar from FAPs cultured under normoxia or hypoxia (n=3). **(C)** Representative photomicrographs showing the morphologies of FACS-sorted satellite cells, PDGFRα+ cells and dermal skin fibroblasts cultured for 7 days and treated for 48h with 5mM 2-DG (n=3). Scale bar, 100μM. **(D)** Percentage of Caspase 3+ cells in satellite cells, FAPs and dermal skin fibroblasts cultured for 5 days and treated for 24h with 5mM 2-DG (n=3). In all graphs values represent the mean ± s.e.m. \**P*<0.05, \*\**P*<0.01 and \*\*\**P*<0.001.

### Inhibition of glycolysis results in the downregulation of the adipogenic/fibrogenic profile and induces endothelial differentiation of FAPs through an mTORC2-HIF2α-eNOS pathway

The mesh network formed by FAPs in response to 2-DG exposure suggested that these cells were moving towards an endothelial cell fate. In contrast to satellite cells, FAPs show low levels of caspase activity (Fig 3D), which indicates that these cells are not undergoing cell death in response to 2-DG, although this marked change in cell morphology could be a result of cell stress. Therefore, we determined mRNA and protein expression levels for key markers of angiogenesis, adipogenesis, and fibrogenesis following 2-DG exposure. Levels of *Lpl* and *Type I collagen* (*Col1a1*) mRNA expression were markedly decreased following 2-DG treatment regardless of oxygen levels (Fig. S3). We noted that commercial media designed for endothelial differentiation had a comparable effect to 2-DG (Fig. S3). Surprisingly, while *Cd31* and *von Willebrand factor (vWF)* transcript levels were unchanged regardless of conditions (Fig. S3). ECs can be identified based upon the binding of *lycopersicon esculentum* lectin to membrane proteins (*37*). We observed that 2-DG and commercial media induced strong lectin binding in FAPs as compared to controls, regardless of oxygen levels (Fig. S4). We conclude that the EC fate is induced by glucose deprivation and is not dependent upon oxygen levels and that the cellular responses to changes in oxygen and glucose levels involve separate signaling pathways. Consistent with this proposal, we observed that FAPs grown under normoxic high glucose conditions accumulated lipid droplets shown by Oil O red staining and Perilipin (Fig. 4A) and expressed Collagen 1 (Fig. 4A). Furthermore, we observed that 2-DG induced, at the protein level, CD31, vWF and ve-cadherin proteins expression and blocked lipid droplet formation and Collagen 1 expression in normoxic FAPs (Fig. 4A). Whereas transcript levels for *vWF* and *Cd31* were unaffected by 2-DG treatment (Fig. S3), we observed an induction of protein expression (Fig. 4A), suggesting that glucose deprivation-induced EC fate involves post-transcriptional regulation.

**Fig. 4.**
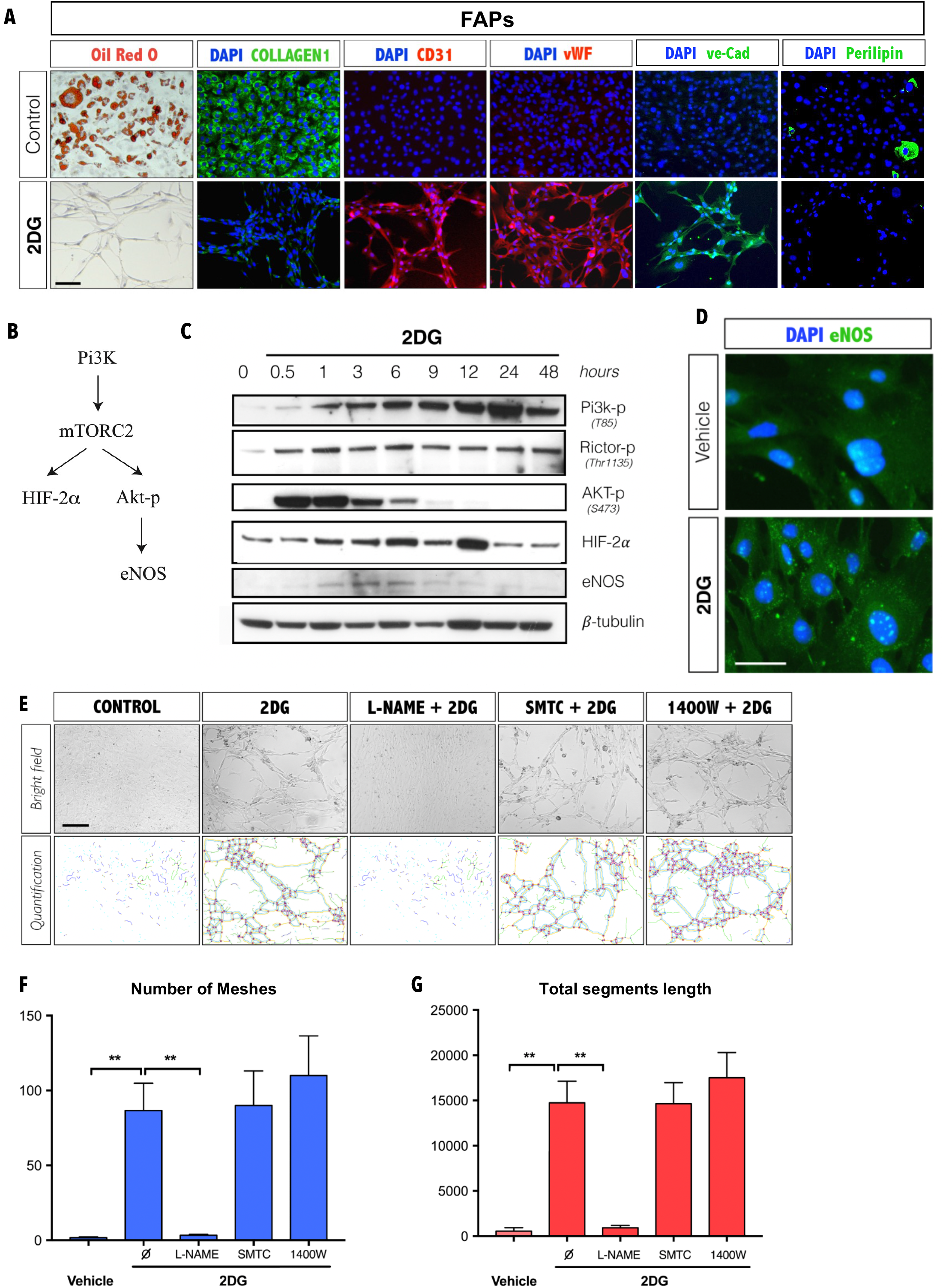
Glycolysis inhibition triggers vasculogenesis of FAPs through an mTORC-2-HIF2α-eNOS pathway. **(A)** Adipogenic (Oil red O and perilipin), fibrogenic (Type I collagen) and vasculogenic (CD31, vWF and ve-cad) cell fates of FAPs (PDGFRα+ cells) following glucose or 2-DG exposure under normoxia (n=3). Scale bar, 100μM. **(B)** Schematic representation of signaling pathway effectors involved in angiogenesis. **(C)** Western-blot from protein extract of FAPs (PDGFRα+ cells) treated with 2-DG at times indicated (n=3). **(D)** Immunostaining of eNOS on FAPs (PDGFRα+ cells) after 6h of 2-DG treated. Nuclei were counterstained with DAPI (n=3). Scale bar, 20μM. **(E)** FAPs (PDGFRα+ cells) were cultured for 48 hours with 2-DG with specific inhibitors for nNOS (SMTC), iNOS (1400W) and pan-NOS inhibitor (L-NAME) as indicated. Scale bar 100μM. Lower panel shows mesh and segment/length images generated from angiogenesis ImageJ (n=3). **(F)** Quantification of the number of meshes using the angiogenesis analyzer of Image J as shown in panel E (n=3). **(G)** Quantification of the total segment length using the angiogenesis analyzer of Image J as shown in panel E (n=3). In all the graphs values represent the mean ± s.e.m. \**P*<0.05, \*\**P*<0.01 and \*\*\**P*<0.001.

Several signaling effectors have been demonstrated to be required for angiogenesis, including Phosphoinositide 3-kinase (Pi3K), RPTOR independent companion of MTOR, complex 2 (Rictor), thymoma viral proto-oncogene 1 (Akt), Hypoxia-inducible-factor (HIF-2α), and endothelial NOS (eNOS) (*38–41*). Previous studies have linked these effectors in a common pathway (*40–43*) (Fig. 4B). Specifically, pi3k-mTORC2 regulates the transcription of HIF-2α and induces angiogenesis through the release of nitric oxide (NO) by the induction of endothelial nitric oxide synthase (eNOS) (*40–42*). In addition, the mTORC2 complex has been shown to activate HIF-2α (also referred to as endothelial PAS domain-containing protein 1) and to induce Akt phosporylation on Serine 473 (s473) that in turn regulates eNOS activity (*40–43*). Western blot analyses of FAPs during the first 48 hours following treatment with 2-DG revealed a rapid induction of Pi3k, Rictor (subunit of mTORC2), and phosphorylated Akt by 1h of 2-DG treatment, followed by an increase in eNOS and HIF-2α (Fig. 4C and Fig. S5). We confirmed eNOS protein expression in cultured FAPs in response to 2-DG (Fig. 4C, D). eNOS is a member of the NOS family that includes neuronal NOS (nNOS, NOS1) and inducible NOS (iNOS, NOS2) (*44–46*). Several inhibitors have been developed to the NOS pathway including a pan-NOS inhibitor N(ω)-nitro-L-arginine methyl ester (L-NAME), as well as an iNOS (1400W) and nNOS inhibitors (S-methyl-L-thiocitrulline (SMTC) (*44–46*), whereas no specific inhibitor of eNOS is currently available. Treatment with the pan-NOS inhibitor (L-NAME) completely blocked 2-DG induced tube/segment formation, whereas inhibition of nNOS and iNOS had no effect (Fig. 4E-G). We conclude that glucose deprivation triggers endothelial differentiation in FAPs through an eNOS dependent pathway and is mediated by effectors known to regulate angiogenesis (*38, 40–42*). We further conclude that while these cells have previously been characterized as fibroadipogenic progenitors, they also constitute an endothelial progenitor population.

### FAPs form capillaries following muscle injury *in vivo*

Our observations that hypoxia and glucose deprivation occur at the site of muscle injury *in vivo* and are also signals that promote the proliferation and subsequent recruitment of FAPs towards the EC fate *in vitro*, suggested that FAPs represent a muscle resident population of vascular progenitors that are enrolled in response to muscle injury. We recently showed that the Osr1^GCE-mTmG^ mouse line can be used to track the FAP lineage cells following injury in postnatal muscle, such that Osr1^GCE-mTmG^ mice can be used to lineage trace the FAP population in skeletal muscle (*47, 48*). This genetic lineage model confirmed that the ectopic fatty infiltration was derived from Osr1 lineage-marked cells following glycerol-induced injury (*48*). Glycerol-induced muscle injury results in poor skeletal myofiber and vascular regeneration (*49*). To determine whether FAPs give rise to vascular tissue *in vivo* during muscle regeneration, TA muscle was injured by both focal freeze-crush and cardiotoxin injury (CTX) in 6 months old Osr1^GCE-mTmG^ mice followed by 5 daily tamoxifen injections and allowed to undergo complete regeneration for 1 month (Fig. 5A). Whereas we did not observe any GFP labeled cells in uninjured muscle, we observed strong GFP expression following injury (Fig. 5B). Using CD31 to identify ECs, we observed that Osr1-GFP cells were distributed into 5 distinct compartments: muscle interstitium, small vessel associated, small vessel co-localized, big vessel associated, and big vessel co-localized (Fig. 5B, C). The majority of Osr1-GFP+ cells were found in the interstitial space (50%; Fig. 5B, C) whereas 20% were found associated with small vessels, 10% associated with big vessels, and very few were in big vessels, however we note that the frequency of big vessels was low in the TA (Fig. 5B, C). Importantly, we observed that ~20% of Osr1-GFP+ cells co-localized with CD31 in small vessels (Fig. 5B, C, D). The Osr1-GFP labeled small vessels are composed of a single layer of CD31+ cells with no obvious closely associated cell types, consistent with their identity as capillaries (Fig. 5D). In order to unequivocally determine the identity of the Osr1-GFP+ cells and confirm their presence in capillaries, we co-stained injured muscle tissue with other EC specific maker, such as Claudin 5 (Cld5, Fig. 5E) and ve-cadherin (ve-Cad, Fig. 5F). Similar results were obtained when the muscle was injured by CTX for the 3 EC specific marker used (CD31, Cld5 and ve-cad, Fig. S6). Taken together, we conclude that FAPs are a highly plastic stromal progenitor cells in skeletal muscle tissue that not only give rise to fat and fibrotic tissue during pathological skeletal muscle regeneration, but also give rise to new vasculature during functional regeneration.

**Fig. 5.**
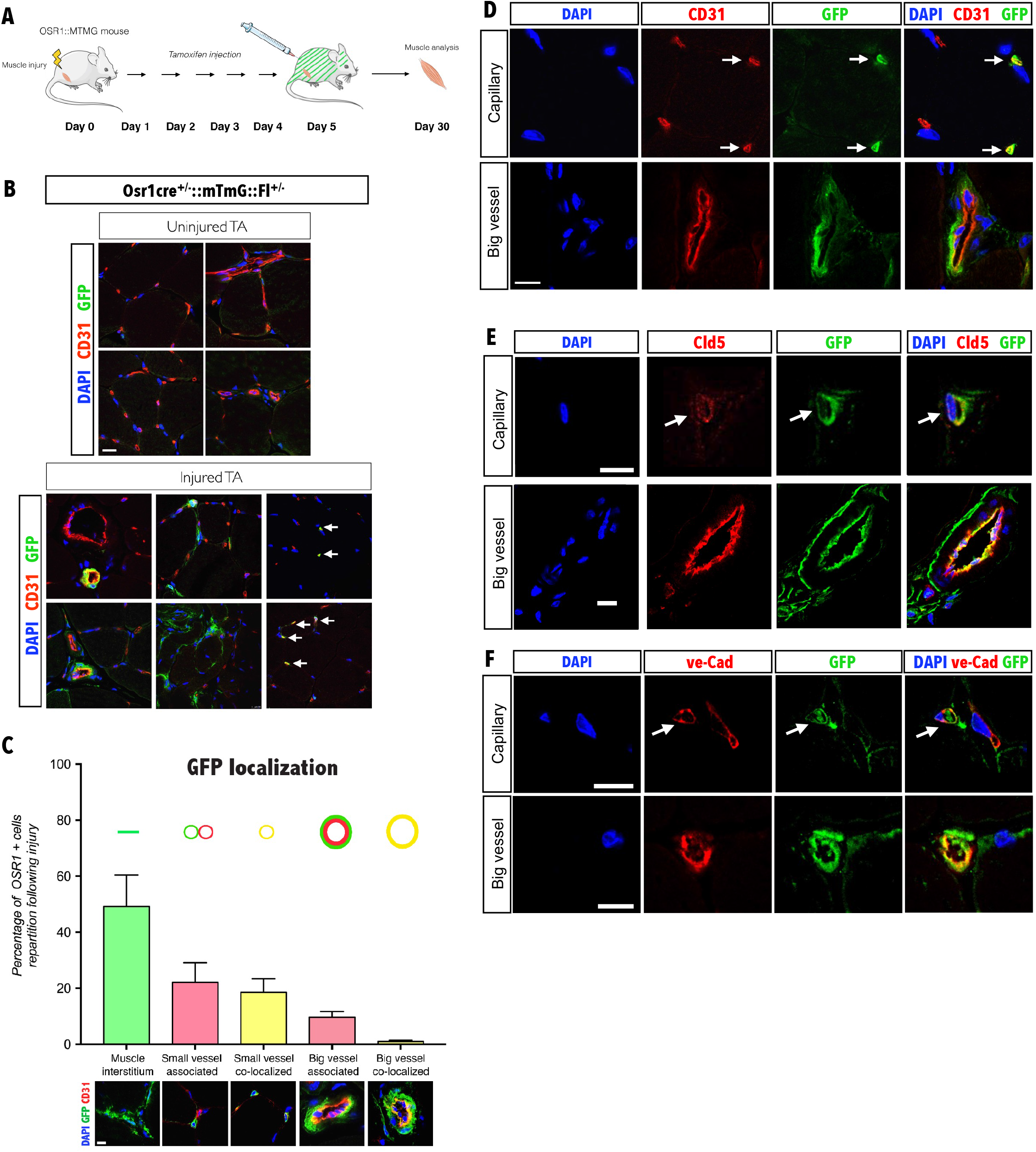
FAPs give rise to capillaries in vivo following muscle injury. **(A)** Osr1:^MTMG^ mouse TA were injured (both freeze-crush and CTX injury) and tamoxifen was injected during the 5 first days following muscle injury and muscles were analyzed one month following injury. **(B)** CD31, GFP and DAPI labeling of uninjured and 1 month injured TA from Osr1:^MTMG^ mice. Arrows denote cells with colocalized GFP (Osr1 lineage positive) and CD31 expression (n=3). Scale bar, 20μm. **(C)** Quantification of the localization of Osr1-GFP+ cells in regenerated TA (n=5). Scale bar, 20μm. **(D)** CD31, GFP and DAPI labeling in regenerated TA from Osr1:^MTMG^ mouse. Arrows denote cells with colocalized expression of GFP (Osr1 lineage positive) and CD31 expression (n=3). **(E)** Claudin 5 (Cld5), GFP and DAPI labeling in regenerated TA from Osr1:^MTMG^ mouse. Arrows denote cells with colocalized expression of GFP (Osr1 lineage positive) and Claudin 5 expression (n=3). **(F)** ve-Cadherin (ve-Cad), GFP and DAPI labeling in regenerated TA from Osr1:^MTMG^ mouse. Arrows denote cells with colocalized expression of GFP (Osr1 lineage positive) and ve-Cadherin expression (n=3). Scale bar, 10μm.

## Discussion

The skeletal muscle interstitium has been proposed to be a site of resident vascular progenitor cells, although the clear identity of these cells remained unresolved and the cell fates obtained *in vitro* appeared stochastic (*50*). Blood vessel-associated vascular progenitor cells have been characterized for large vessel formation (*51, 52*), however the identity of endothelial progenitors and signals that direct the repair the microvasculature in response to injury are poorly understood. We propose that FAPs act as a stress sensitive stromal progenitor population that responds to a decrease in oxygen and glucose levels by proliferation and endothelial differentiation, respectively. Consistent with this proposal, we observed that the number of FAPs (PDGFRα+/CD31- cells) increased at 3 and 5 dpi, whereas ECs (CD31+PDGFRα- cells) emerged by 7 and 14 dpi. Previous studies implicated the HIF-1α-mTORC1 pathway in proliferation, as well as the HIF-2α-mTORC2 axis that act to promote cell survival and angiogenesis (*53–55*). Accordingly, we found that hypoxia induces proliferation of FAPs through mTORC1-HIF1α-Glut1 pathway, whereas 2-DG induces endothelial differentiation via mTORC2-HIF-2α-eNOS signaling (Fig. 6). Our data are consistent with a recent report demonstrating that glutamine withdrawal enhances ECs differentiation in human embryonic stem cells (*56*), suggesting that perturbation of the energy metabolism promotes angiogenesis. Skeletal muscle contains resident progenitor cells able to differentiate into adipocytes upon high glucose exposure, while hypoxia had been previously suggested to inhibit adipogenesis (*57, 58*). Here we show that hypoxia inhibits adipogenesis in FAPs, even in the presence of high concentrations of glucose, and that glucose deprivation induces FAPs to downregulate both fibrotic and adipogenic profiles and upregulate the EC program, regardless of oxygen levels. Thus, the ability of FAPs to proliferate and differentiate into adipocytes, fibroblasts or endothelial cells is tightly regulated by oxygen and glucose availability. The metabolic switch controlling FAP cell fate may contribute to the vascular complications that occur in diseases such as diabetes or tumor growth (*59*). Our results point to potential therapeutic interventions in vascular disease and in promoting vascular regeneration.

**Fig. 6.**
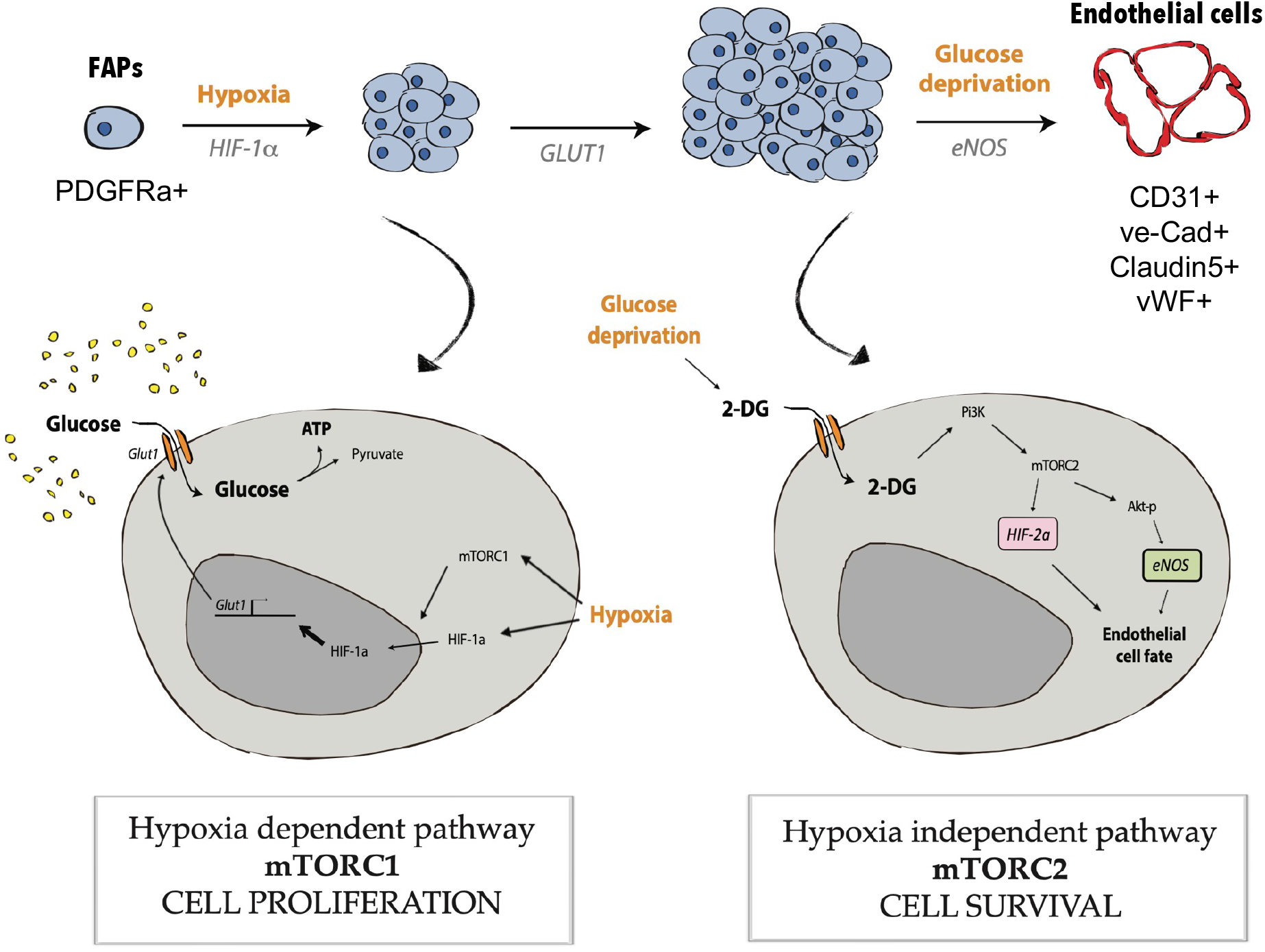
Summary schema showing pathways leading to FAPs proliferation, survival and endothelial cell fate. The injured muscle becomes quickly hypoxic, inducing the proliferation of FAPs through an mTORC1-HIF1a-Glut1 pathway. The proliferation of glycolytic cells under a non-vascularized hypoxic tissue leads to glucose deprivation, which induces endothelial differentiation of FAPs through an mTORC2-HIF-2a-eNOS pathway.

## Acknowledgment

We thank the flow cytometry platforms (Catherine Blanc from CyPS Pierre et Marie Curie University and to Camille Knosp from PARCC), to the animal facility platform for their help handling mouse lines (to the Pierre et Marie Curie University animal facility and to ERI at PARCC).

This work received funding from the European Community Seventh Framework Programme project ENDOSTEM (Activation of vasculature associated stem cells and muscle stem cells for the repair and maintenance of muscle tissue) (Agreement 241440), Laboratoire d’Excellence Revive (Investissement d’Avenir, ANR-10-LABX-73), The Fondation Leducq (grant 13CVD01; CardioStemNet project) and the Agence Nationale pour la Recherche (ANR) grant RHU CARMMA (ANR-15-RHUS-0003). In addition, this work was funded by the German Research Foundation (DFG; grant GK1631), French-German University (UFA-DFH; grant CDFA-06-11), the Association Française contre les Myopathies (AFM 16826), and the Fondation pour la Recherche Médicale (FRM DEQ20140329500). Both teams are beneficiaries of the MyoGrad International Research Training Group for Myology.

## Author contributions

D.O. designed and carried out most experiments, analysed and interpreted the data and wrote manuscript. V.B. assisted with in vitro and in vivo Osr1 mouse line experiments. R-M.C. assisted with satellite isolation and cultures. A.S. assisted with in vitro experiments. B.H. performed FACS. S.P.K. assisted with in vivo Osr1 mouse line experiments. S.S. and J.S.H. assisted with critical feedback on the manuscript. M.V. performed FACS, in vivo Osr1 experiments, assisted with analysis and critical feedback on the manuscript. G.M. and D.S. conceived and supervision of the project and manuscript, and secured funding.

## MATERIAL AND METHODS

### Mice

Satellite cells and FAPs were FACS-sorted from 3 month-old C57BL/6JRj mice (Janvier) for *in vitro* analyses. Lineage tracing experiments were performed on 6 month-old Osr1^GCE/+^:R26^mTmG/+^ mice(*60*) to track FAPs (*47, 48*). Lineage tracing of injury-activated Osr1-GFP^+^ FAPs was performed by intraperitoneal injection of 3 mg of Tamoxifen starting at the day of injury and continued for a total of 5 days. All work with mice was carried out in adherence to national and European guidelines (CEEA34 and French ministry of research and Directive 2010/63/EU).

### Fluorescence-activated cell sorting analysis

For fluorescence-activated cell sorting (FACS), limb muscle were minced and digested in Hank’s Balanced Salt Solution (HBSS, GIBCO) containing 2 μg ml^−1^ collagenase A (Roche), 2.4 U ml^−1^ dispase I (Roche), 10 ng ml^−1^ DNase I (Roche), 0.4 mM CaCl and 5 mM MgCl as described previously(*1, 2*). Primary antibodies used at a concentration of 10 ng ml^−1^ were: rat anti-mouse CD45-PeCy7 (eBiosciences), rat anti-mouse TER119-PeCy7 (BD Biosciences), rat anti-mouse CD34-BV421 (BD Biosciences), rat anti-mouse SCA1-FITC (eBiosciences), rat anti-mouse PDGFRα-PE (eBiosciences), rat anti-mouse CD31-BUV737 (BD Biosciences) rat anti-mouse α7INTEGRIN-A700 (RD Biosystem) and rat anti-mouse CD11b-BUV395 (BD Biosciences). Cells were incubated for 30 minutes on ice and washed once with ice-cold HBSS, filtered and re-suspended in HBSS containing 0.2% (w/v) bovine serum albumin (BSA), 1% (v/v) penicillin-streptomycin and 10 ng ml^−1^ DNase I. Cells were incubated with the viability marker 7-AAD (BD Bioscience) for 10 minutes at 4°C prior to cells sorting. Flow cytometry analysis and cell sorting were performed on a FACS Aria (Becton Dickinson) with appropriate Fluorescence Minus One controls. FAPs were negatively selected for TER119, CD45, CD11b, α7INTEGRIN, and CD31 and purified based on the expression of CD34, SCA1, and PDGFRα. Satellite cells were negatively selected for TER119, CD45, CD31, PDGFRα and SCA1 and purified based on the expression of α7INTEGRIN and CD34 (Fig. S1).

### Primary cell culture

Freshly FACs purified cells were plated on gelatin-coated dishes at a density of 3000 cells per cm^2^. Cells were grown for one week in amplification medium (AM) containing high-glucose (4.5 g/l) Dulbecco’s modified Eagle medium (DMEM; Gibco) supplemented with 20% heat-inactivated fetal bovine serum (FBS; Hyclone), 10% heat-inactivated horse serum (Gibco), 1% (v/v) penicillin-streptomycin (Gibco), 1% (v/v) L-glutamine (Gibco), and 1% (v/v) sodium pyruvate (Gibco). For adipogenic differentiation, cells were cultured for 10 days in AM. Cells were cultured under normoxic conditions (21% O_2_) or hypoxic conditions (1.5% O_2_) using a hypoxia incubator chamber (Stem Cells Technologies) and oxygen levels were monitored with an oxygen sensor probe to follow oxygen tension in real-time (GasBadge Pro., Industrial Scientific). For endothelial differentiation, cells were placed for 2 days in Endothelial Cell Growth Medium 2 (EGM-2; PromoCell) supplemented with 0.5 ng/ml of vascular endothelial growth factor 165 (VEGF), and 1% (v/v) penicillin-streptomycin. We note that EGM-2 contains 1g of glucose/l as per manufacter’s information. The media was changed every 2 days.

### Cell proliferation assay

Cell proliferation was performed by using the CyQuant^®^ NF Cell Proliferation Assay Kit (Invitrogen) on freshly purified FAPs. FAPs were seeded at 1250/cm22 in a 96 well-plate and cultured in normoxic condition (37°C, 21% O22) or hypoxic condition (37°C, 1% O22) as mentioned above. Cells were incubated with 100μL of dye binding solution during 1h at 37°C according to the manufacturer’s instructions after 4 or 6 days in culture. The fluorescence intensity was measured at ambient temperature using the fluorescence microplane reader (Flex Station 3, Molecular Devices) with excitation at 485nm and emission detection at 530nm.

### Pharmacological inhibitors

Cells were FACS-sorted and cultured in AM for 7 days with 10μM of Wzb-117 or with 100 nM Rapamycin (Sigma). For glucose deprivation experiment, cells were grown in AM for 7 days, and exposed to 5 mM of 2-deoxy-D-glucose (2-DG) at time intervals indicated. Cells were pretreated for 24 hours with different NOS inhibitors: L-NAME (Sigma, 2 mM), SMTC (Sigma, 1mM) or 1400W (Sigma, 1mM) and treated with 2-DG for 48h. All inhibitors were resuspended in DMSO.

### RNA interference

Freshly sorted cells were expanded and passaged 1 time (P1) and then seeded at 500 cells/cm^2^ and transfected for 24 hours with 20 nM of siRNAs resuspended in Opti-MEM (Gibco), DharmaFECT siRNA transfection reagent (Fisher scientific) without antibiotics. The medium was changed every 2 days with AM and cell proliferation was analyzed by counting the number of crystal violet (Sigma) positive cells. siRNA scrambled: (Ambion), siRNA HIF-1α (target sequence: CACCATGATATGTTTACTA, Eurogentec).

### RNA extraction and qRT-PCR

Total RNA extracts were prepared from a minimum of 2×10^4^ freshly sorted cells using RNeasy Micro Kit (Qiagen) according to manufacturer’s instructions, and reverse transcribed using the SuperScript II First-Strand Synthesis System (Life Technologies). Semi-quantitative PCR was performed using ReddyMix Master Mix (Thermo Scientific) under the following cycling conditions: 94°C for 5 minutes followed by 30 cycles of amplification (94°C for 30 seconds, 60°C for 30 seconds and 72°C for 1 minute) and a final incubation at 72°C for 10 minutes.

### Real time PCR primers

*Hif-1a* forward TCAAGTCAGCAACGTGGAAG, *Hif-1a* Reverse TATCGAGGCTGTGTCGACTG. *Glut1* forward GCTCTACGTGGAGCCCTA, *Glut1* reverse CACATCGGCTGTCCCTCGA. *Lpl forward* CTGCTGGCGTAGCAGGAAGT, *Lpl reverse* GCTGGAAAGTGCCTCCATTG. *Pparδforward* CAAGAATACCAAAGTGCGATCAA, *Pparδreverse* GAGCAGGGTCTTTTCAGAATAATAAG. *Acr30 forward* GCTCCTGCTTTGGTCCCTCCAC, *Acr30 reverse* GCCCTTCAGCTCCTGTCATTCC. *Col1a1 forward* GCTCCTCTTAGGGGCCACT, *Col1a1 reverse* CCACGTCTCACCATTGGGG. *Cd31 forward* GAGCCCAATCACGTTTCAGTTT, *Cd31 reverse* TCCTTCCTGCTTCTTGCTAGCT*. vWF forward* CCGGAAGCGACCCTCAGA, *vWF reverse* CGGTCAATTTTGCCAAAGATCT. *Rpl13 forward* ACTCTGGAGGAGAAACGGAAGG, *Rpl13 reverse*, CAGGCATGAGGCAAACAGTC.

### Lactate production measurements

FAPs were freshly isolated and cultured for 7 days in AM in 48-well plates and exposed to normoxic (21% O_2_) or hypoxic (1.5 % O_2_) conditions with serum-free amplification medium for 16 hours. Supernatants were recovered and lactate content was analyzed using a lactate colorimetric assay kit (K607, BioVision, Milpitas, USA). Cells were lysed with RIPA and lactate concentration was normalized to protein content.

### Western blot

Cells were homogenized in lysis buffer (150mM NaCl, 50mM Hepes pH7.6, 1% NP-40, 0,5% sodium deoxycholate, 5mM EDTA) supplemented with 1 mM PMSF, Complete (Roche), 20mM NaF, 10mM b-glycerophosphate, 5mM Na-pyrophosphate, and 1mM Na-orthovanadate. Protein concentration was measured using the Pierce^®^ BCA protein Assay kit (Thermo Scientific) and separated by electrophoresis (Novex NuPAGE Bis-Tris protein gel 4–12% or 5% home-made Bis-Tris gel) and transferred to a PVDF membrane in 20% methanol transfer buffer. Membranes were probed with anti-Pi3K (Cell Signaling), anti-Akt-p (Cell Signaling), anti-Rictor-p (Cell Signaling), anti-HIF-2α (Abcam), anti-eNOS (Thermo fisher) and anti-α-tubulin (Sigma). Antibody binding was visualized using horse-radish peroxidase (HRP)-conjugated species-specific secondary antibodies (Jackson ImmunoResearch) followed by enhanced chemiluminescence (Pierce).

### Histological and cell analyses

Cryosections (10μm) were prepared from tibialis anterior snap frozen in liquid nitrogen-cooled isopentane. Tissue sections, cultured cells, and cytospin preparations were fixed in 4% (w/v) paraformaldehyde and processed for immunostaining as described previously(*2*). Primary antibodies used were: anti-HIF-1α (Active motif), anti-CD31 (BD bioscience), anti-vWF (Abcam), anti-ve-cadherin (R&D), anti-claudin 5 (Thermofisher), anti-laminin (Sigma), anti-GFP (Sigma), anti-Collagen 1 (Abcam), anti-caspase3 (BD bioscience), eNOS (Abcam) and anti-Perilipin (Abcam). Antibody binding was revealed using species-specific secondary antibodies coupled to Alexa Fluor 488 (Molecular Probes), Cy3 or Cy5 (Jackson ImmunoResearch). Nuclei were counterstained with DAPI (Sigma). Live cells were incubated with 20 μg/ml *Lycopersicon esculentum lectin* (Dylight 594, Vector laboratories) for 30 minutes at 37°C and washed with PBS. Images were acquired using a Leica DM-IL inverted fluorescence and light microscope, Leica DM fluorescence and light microscope or Leica SPE confocal microscope. To stain lipids, cells were fixed in 10% formalin (Sigma) for 5 minutes at 4°C, rinsed in water and then incubated in 100% propylene glycol (Sigma) for 10 minutes, stained with Oil Red O (Sigma) for 10 minutes at 60°C, placed in 85% propylene glycol for 2 minutes and rinsed in water. Nuclei were counterstained with Mayer’s Hematoxylin Solution (Sigma). To analyze cell proliferation, cells were stained with 0.5 % (m/v) crystal violet (Sigma) for 30 minutes at room.

### Regeneration assays

Skeletal muscle regeneration was induced by intramuscular CTX injection (0.06mg/ml, Sigma) and muscles were analyzed 1, 4, 5, 6, 7 and 14 days after injury. To analyze OSR1-GFP+ cell fate from Osr1^GCE/+^:R26^mTmG/+^ mice, muscle regeneration was analyzed 1 month following focal freeze crush injury, as previously described (*48*), or CTX injury.

### Pimonidazole injection and detection

Mice were injected (Intraperitoneal) with 60 mg/kg of Pimonidazole and muscle was harvested 90 minutes following injection and placed in liquid nitrogen-cooled isopentane. Cryosections (10 μm) of TA muscles were fixed in 4% (w/v) paraformaldehyde. Pimonidazole was detected using the Hypoxyprobe-1™ kit (Hydroxyprobe,Burlington, USA) according to manufacturer’s instructions. Antibody binding was revealed using species-specific secondary antibody conjugated to Cy3 (Jackson ImmunoResearch),

### 2-NBDG injection and detection

The fluorescent glucose analogue probe (2-NBDG) was injected (Retro-orbital, RO) at a concentration of 16 mg/kg 5 minutes before harvesting muscle. Muscle tissue was placed immediately into liquid nitrogen-cooled isopentane. Cryosections (10 μm) were analyzed using a Leica DM-fluorescent microscope.

### Statistical and data analyses

All statistical analyses were performed using an unpaired Student’s *t*-test in the StatView software. Values represent the mean ± s.e.m. \**P*<0.05, \*\**P*<0.01 and \*\*\**P*<0.001. All histological and cell experiments were repeated at least 3 times in independent experiments and representative results are shown.

---

**Fig. S1:**
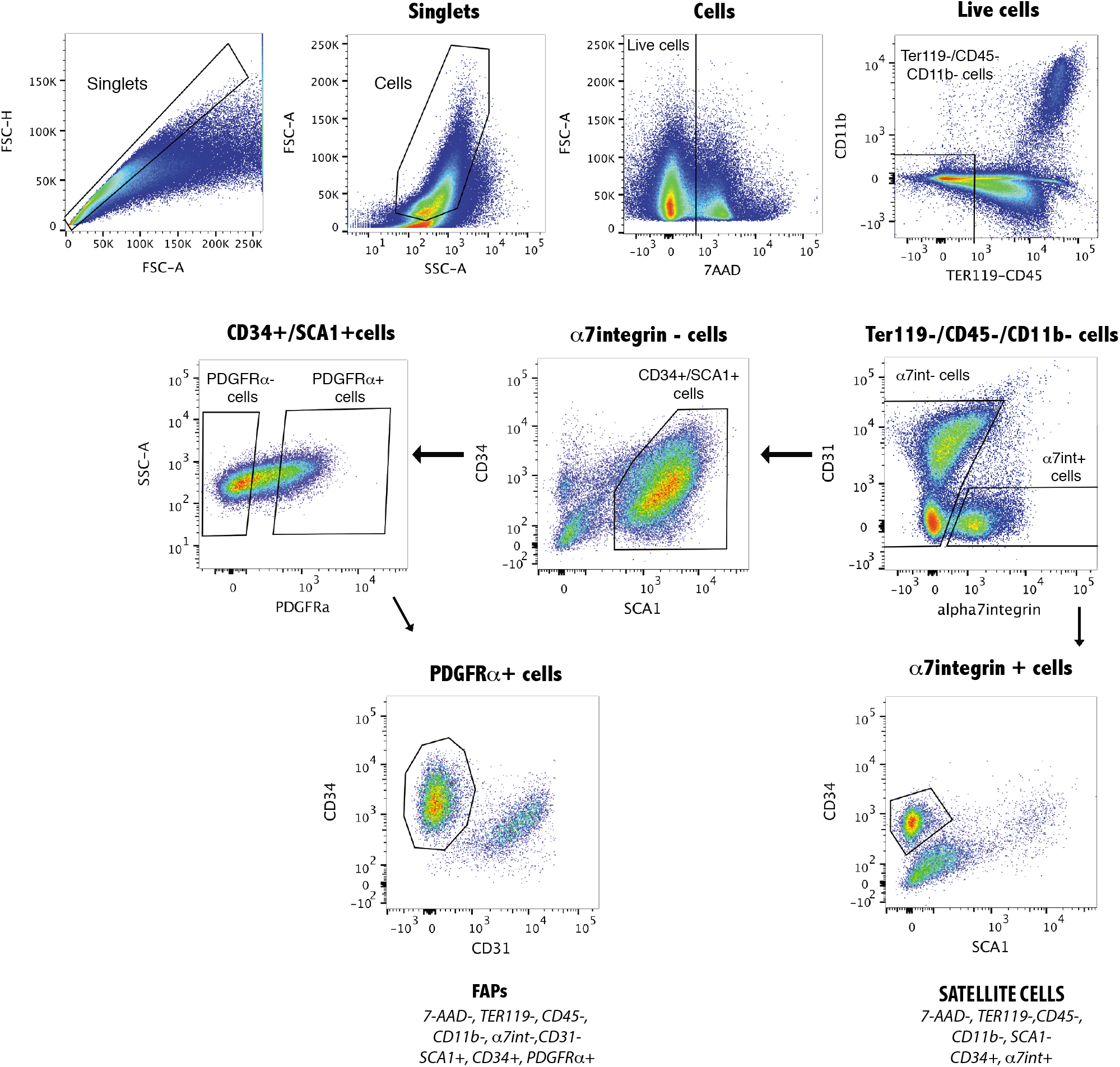
Cell sorting strategy for FAPs and satellite cells isolation. A, Forelegs muscle cell suspension were analyzed using the following strategy: live cells were selected with the viability marker 7-AAD (7-AAD negative fraction); Muscle resident cells were selected by exclusion of the hemopoietic cells (CD45, TER119 and CD11b negative fraction); Satellite cells were selected by expression followed by CD34 expression and exclusion of Sca-1; FAPs were selected by exclusion of α7integrin and CD31, followed by expression of CD34 and Sca-1 and finally of PDGFRα expression.

**Figure S2:**
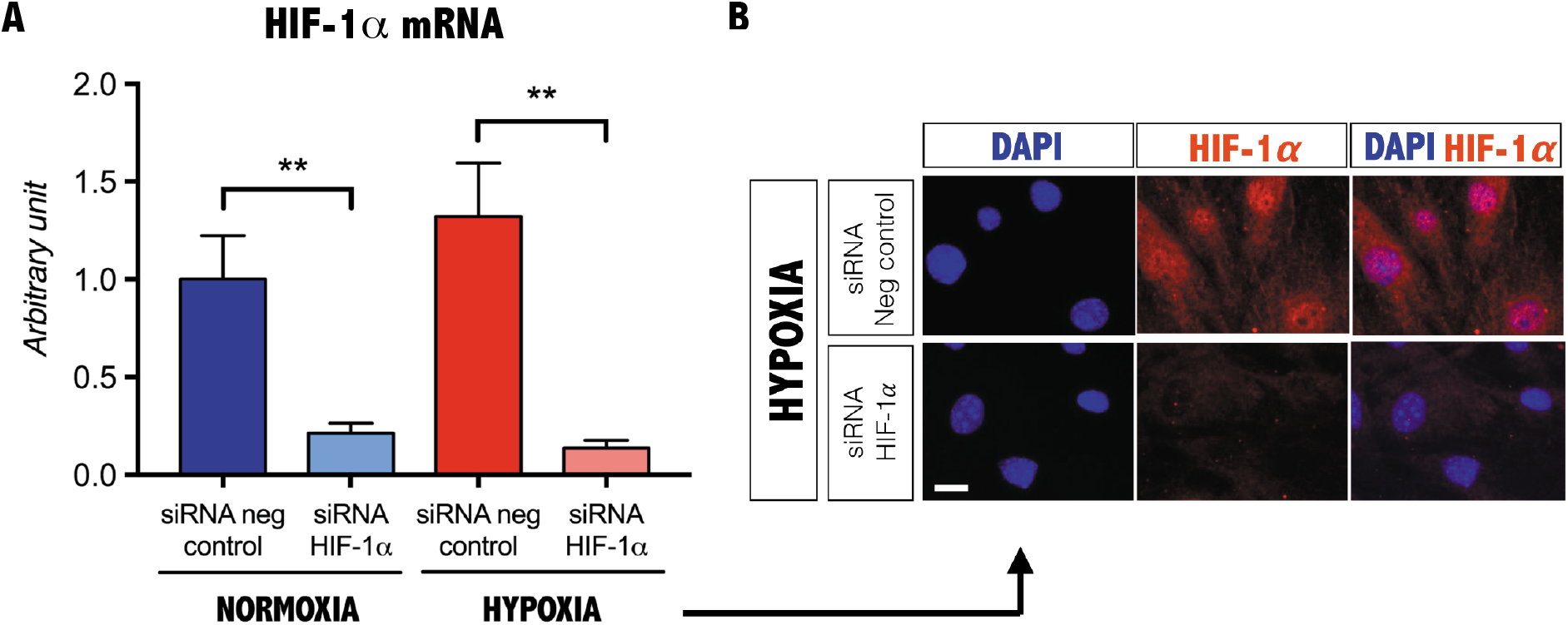
In vitro HIF-1a silencing. A, qPCR of HIF-1α from FAPs cultured under normoxic or hypoxic conditions following siRNA mediated knockdown of HIF-1α (n=3). Values represent the mean ± s.e.m. \**P*<0.05, \*\**P*<0.01 and \*\*\**P*<0.001. B, Immunostaining of HIF-1α following siRNA mediated knockdown of HIF-1α on PDGFRα+ cells cultured under hypoxic conditions (n=3). Scale bar, 10 mm.

**Figure S3:**
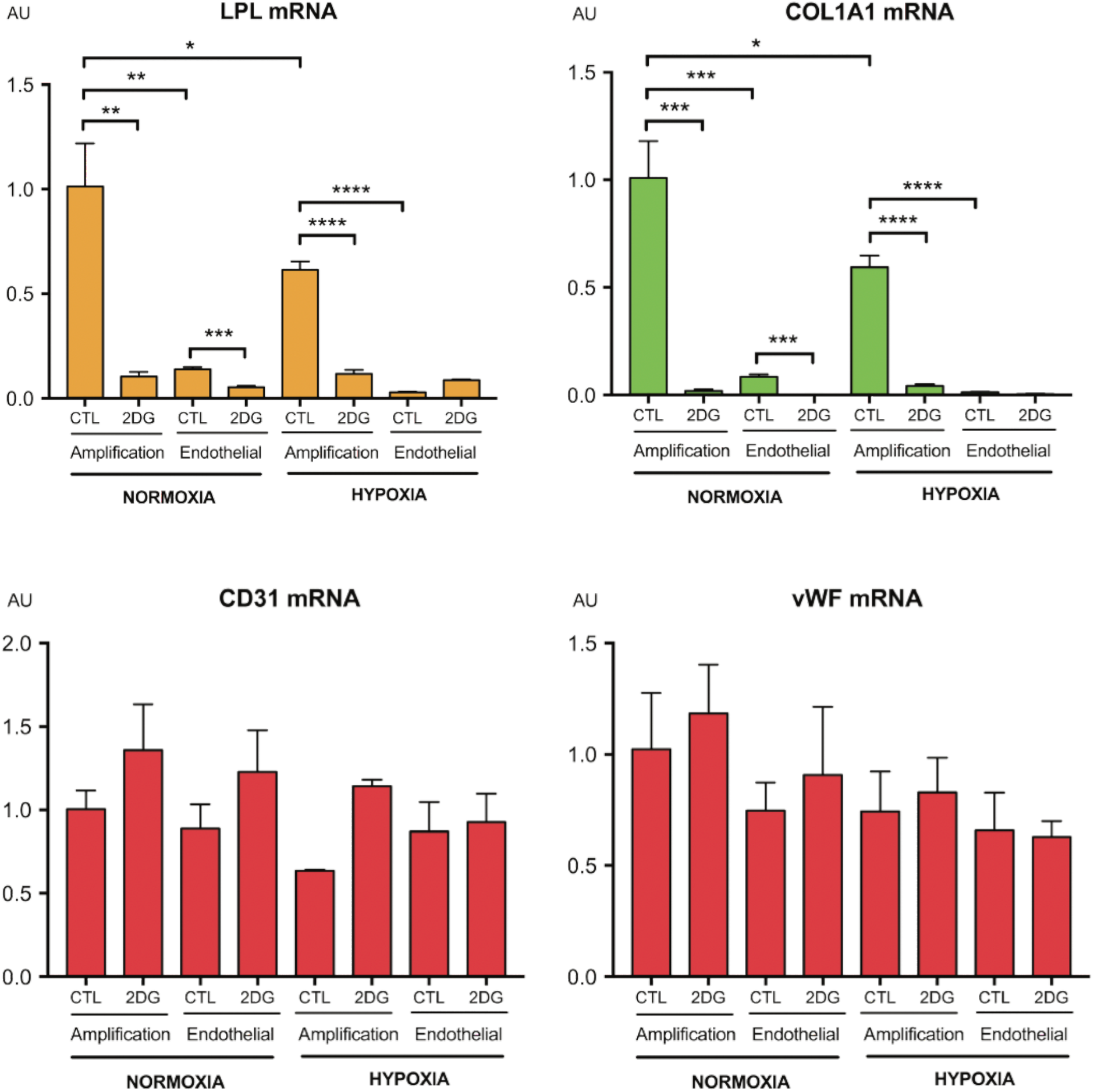
Transcriptional profile of FAPs after in vitro treatment with 2DG and endothelial media. qPCR of *Lpl*, *Col1a1*, *Cd31* and *vWF* from FAPs (PDGFRα+ cells) cultured 48h under normoxic or hypoxic conditions in amplification or endothelial cell medium with or without 2-DG (n=4).

**Figure S4:**
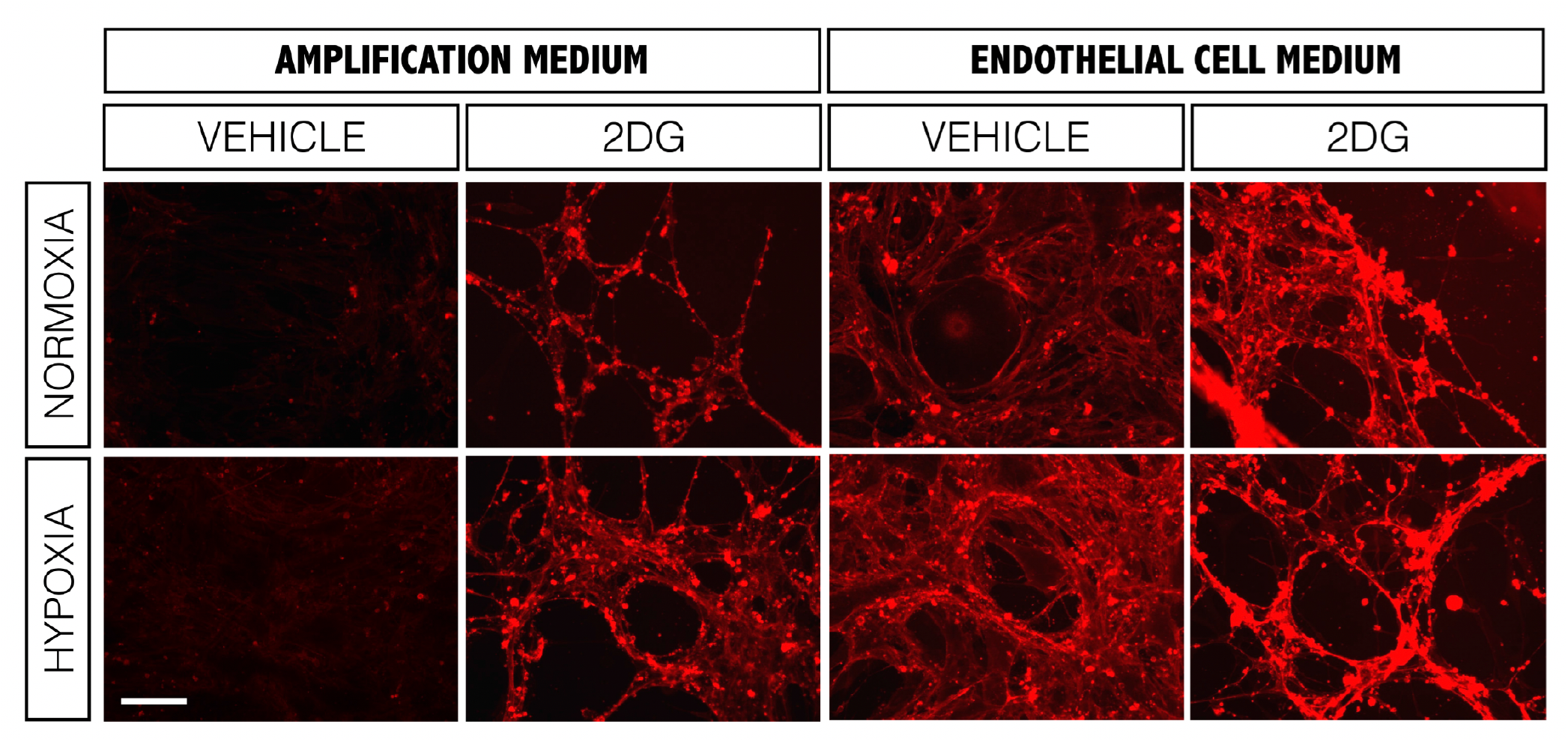
Effect of 2-DG, oxygen and the culture media on lectin binding in FAPs. FAPs were FACS-sorted, cultured for one week high glucose DMEM or low glucose endothelial medium and then treated for 48 hours with 2-DG under normoxia or hypoxia. Living cells were incubated with lectin (20μg/ml) for 30 minutes at 37°C (n=3). Scale bar, 100μm.

**Figure S5:**
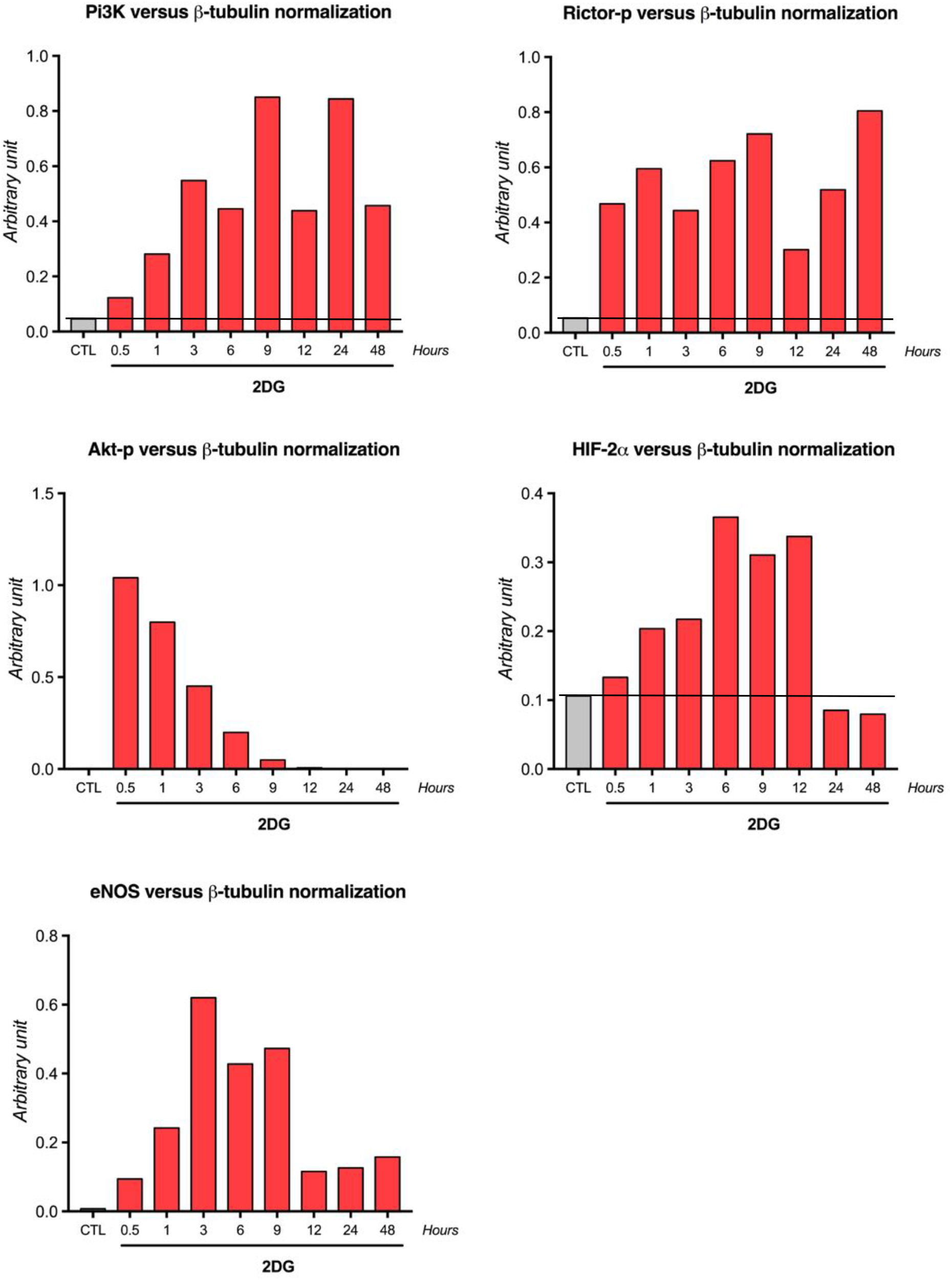
Quantification of the western-blot. PI3K, Akt-p, Rictor-p, HIF-2a, eNOS and b-tubulin bands were quantified and the PI3K, Akt-p, Rictor-p, HIF-2a and eNOS protein expression were normalized to b-tubulin (n=3).

**Figure S6:**
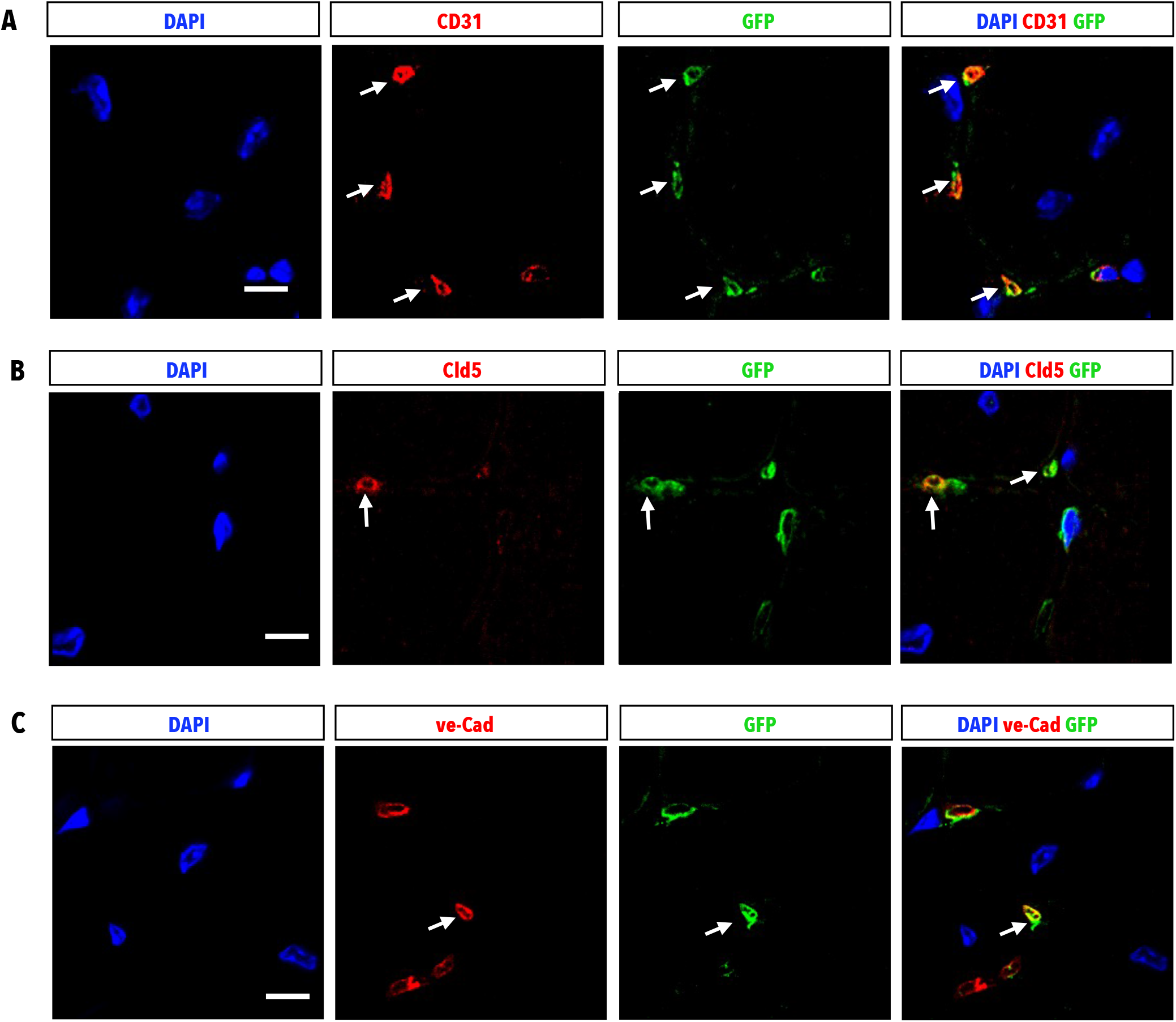
FAPs give rise to capillaries in vivo following cardiotoxin (CTX) injury. A, CD31, GFP and DAPI labeling in regenerated TA from Osr1:^MTMG^ mouse. Arrows denote cells with colocalized expression of GFP (Osr1 lineage positive) and CD31 expression (n=3). B, Claudin 5 (Cld5), GFP and DAPI labeling in regenerated TA from Osr1:^MTMG^ mouse. Arrows denote cells with colocalized expression of GFP (Osr1 lineage positive) and Claudin 5 expression (n=3). C, ve-Cadherin (ve-Cad), GFP and DAPI labeling in regenerated TA from Osr1:^MTMG^ mouse. Arrows denote cells with colocalized expression of GFP (Osr1 lineage positive) and ve-Cadherin expression (n=3). Scale bar, 10μm.

